# Mycorrhizal inoculation and treatment with biochar and compost from pruning waste improve the qualitative properties of a calcareous soil under wheat cultivation

**DOI:** 10.1101/2021.08.14.456348

**Authors:** Roghayeh Vahedi, MirHassan Rasouli-Sadaghiani, Mohsen Barin, Ramesh Raju Vetukuri

## Abstract

Most calcareous soils have relatively low levels of organic matter. To address this issue and improve the qualitative properties of calcareous soils, soils can be treated with mycorrhizal fungi and/or exogenous organic material such as biochar or compost derived from tree pruning waste. To evaluate the effect of pruning waste biochar (PWB) and pruning waste compost (PWC) derived from apple and grape trees combined with arbuscular mycorrhizal fungi (AMF) on the biological indices of calcareous soils, a rhizobox study on wheat plants using a completely randomized design was conducted under greenhouse conditions. The studied factors included the source of the type of organic material applied (PWB, PWC, and control), the nature of the microbial inoculation (inoculation with AMF or no inoculation), and the zone to which the treatments were applied (rhizosphere and non-rhizosphere soil). At the end of the plant growth period, organic carbon (OC), microbial biomass carbon (MBC), microbial biomass phosphorous (MBP), microbial respiration (BR), substrate-induced respiration (SIR), alkaline (ALP), acid (ACP) phosphatase enzyme activities in the rhizosphere and non-rhizosphere soils, and root mycorrhizal colonization were determined. Simultaneous application of a source of organic matter and AMF inoculation significantly increased the OC and biological indices of soil relative to those observed when applying organic matter without AMF inoculation. Additionally, MBC, MBP, ACP and ALP - enzymes activities in the rhizosphere zone were significantly higher than in the non-rhizosphere. AMF increased BR and SIR levels in the rhizosphere by 13.06% and 7.95% compared to non-rhizosphere, respectively. It can be concluded that in calcareous soils with low organic carbon contents, organic amendments such as PWC and PWB can improve soil biological properties by increasing microbial activity and changing the properties of the rhizosphere.

## 1. Introduction

The arid and semi-arid regions have alkaline calcareous soils with low contents of organic matter, which adversely affects crop yields in these regions [1]. One of the most important ways to improve the organic matter content is to properly manage the use of plant residues such as waste material from the pruning of fruit trees. Tree pruning waste can be converted into biochar or compost, which can then be applied to soils to replenish or increase their nutrient content. This improves the soil’s physical and chemical characteristics and is important for supporting the life of soil microorganisms and maintaining a sustainable dynamic equilibrium between the biotic and abiotic components of the soils [2, 3]. Biochar is a carbon-enriched solid generated by the pyrolysis of biomass under conditions with little or no oxygen [4]. The composition and properties of biochar depend on the pyrolysis conditions including the temperature, heating rate, duration, pressure, and input material [5]. Conversely, compost is formed by the decomposition of organic matter mediated by microorganisms under hot, humid, aerobic conditions [6]. Compost is rich in mineral nutrients, some of which are released and made available to plants in soil gradually and continuously [7]. In addition to its physical and chemical properties, soil quality is closely related to its biological characteristics [8]. Biological indicators are thus key descriptors of soil quality, and variation in soil performance should be measured using biochemical parameters and indices (e.g., BR, SIR, MBC, MBP, ACP, and ALP) that represent the diversity of soil microorganisms as well as their distribution and metabolic activity. Soil microbial respiration is essentially a cellular process that involves several biochemical reactions; the rate of microbial respiration is an indicator of both the status and activity of soil microbes but also reflects the rate of organic matter decomposition and cycling of certain nutrients within the soil [9].

Enzymatic activity and microbial biomass are the main biological indicators of soil. It seems that among the biochemical parameters, MBC (41%) and the activity of phosphatase (ACP or ALP) enzymes (28%) have been widely used [10].

There are contradictory reports about the impact of biochar on soil quality. Liu et al. reported that biochar did not significantly influence soil respiration [11]. However, other studies concluded that the pores of biochar provide protective habitats for microorganisms, leading to increases in the soil’s content of various mineral nutrients, energy, and carbon; as a result, biochar administration was found to increase soil biological activity and help maintain soil quality [12, 13]. Another study found that biochar increased microbial biomass and activity and also, it can enhance enzymatic activities in soil; the increase in the activity of enzymes like phosphatases and suggested that the increased activity of microorganisms following biochar administration might improve the availability of nutrients to other microorganisms as a result of increased root exudation [14].

Studies on compost have shown that it contains a large population of microorganisms. Therefore, in addition to increasing the organic matter and nutrient content of the soil, compost administration increases and changes its microbial population [15]. Some rhizosphere microorganisms such as AMF can stimulate soil microbial activity and promote increases in activity of phosphatase and soil microbial biomass [16]. Accordingly, multiple studies have found that treatment with biochar and compost promote mycorrhizal root colonization, spore production, and hyphal expansion, thereby improving soil quality [17, 18]. Active root systems continuously release organic compounds into the rhizosphere that promote the growth and activity of the microbial community in the soil as well as the overall health of the soil system [19, 20].

The effect of compost and biochar is not confined to the placement of organic matter in and outside the rhizosphere; it also strongly affects microbial activity in the soil outside the rhizosphere [21]. When studying rhizosphere processes, it is helpful to confine root growth to a fixed volume of soil in order to increase root density and accelerate the sampling of the rhizosphere soil. To this end, the concept of the rhizobox was developed; a rhizobox is like a pot except that it hinders direct contact between the roots and the soil without disrupting the mobilization of the soil solution [22, 23].

This work investigates the effects of applying biochar and pruning waste compost derived from apple trees and grapevines on some biological characteristics of calcareous soils in the presence of AMF. The combined effects of treatment with biochar and AMF on soil biological processes have not been studied extensively, and most biochar administration studies have focused on acidic soils in tropical and humid climates; consequently, there is little data on their effects in warm and arid regions with non-acidic soils.

## 2. Materials and Methods

### 2.1. Soil sampling and preparation

This study used a randomized factorial complete block design with three replicates. The experiments were conducted under greenhouse condition in rhizoboxes. The factors were the organic matter source (pruning waste biochar [PWB], pruning waste compost [PWC], and control [no added organic matter]), microbial inoculation (AMF and non-inoculation [-AMF]), and the zone to which the treatment was applied (rhizosphere or non-rhizosphere soil). Soil samples were collected from the surface layer (0–30 cm) of a region of non-arable land in Salmas, West Azerbaijan Province, Iran. The samples were air-dried and sieved to pass through a 2 mm mesh, then sterilized in an autoclave at 121°C at 1.5 atm for 2 hours. Before sterilization, some of the soil’s physicochemical properties were determined, as shown in Table 1 [24].

**Table 1.** Physicochemical properties of the studied soil

| soil texture | pH | EC<br>(dS m<sup>-1</sup>) | O.C<br>(%) | CaCO<sub>3</sub><br>(%) | N<br>(%) | P<br>(mg kg<sup>-1</sup>) | K<br>(mg kg<sup>-1</sup>) |
| --- | --- | --- | --- | --- | --- | --- | --- |
| Loamy sand | 7.53 | 0.47 | 0.25 | 14.25 | 0.08 | 7.64 | 98 |

### 2.2. Preparation of biochar and compost

To prepare biochar, fruit tree pruning waste was collected from orchards in Urmia County in West Azerbaijan Province. The waste was then oven-dried at 65 °C for 48 hours. The dried samples were placed in a reactor (a steel cylinder with a diameter of 7 cm and a height of 31 cm) and then heated to 350 °C in an electric furnace to form biochar. The PWC was taken from the research greenhouse of the Soil Science Department in Urmia University. Finally, the PWB and PWC were ground and screened with a 0.5-mm mesh. No ash was observed on the biochar surface, implying that oxygen had been removed and it had been produced correctly. Table 2 summarizes the characteristics of the biochar and compost, which were determined using the analytical methods described by Rajkovich et al. and Alfano et al. [25, 26].

**Table 2.**
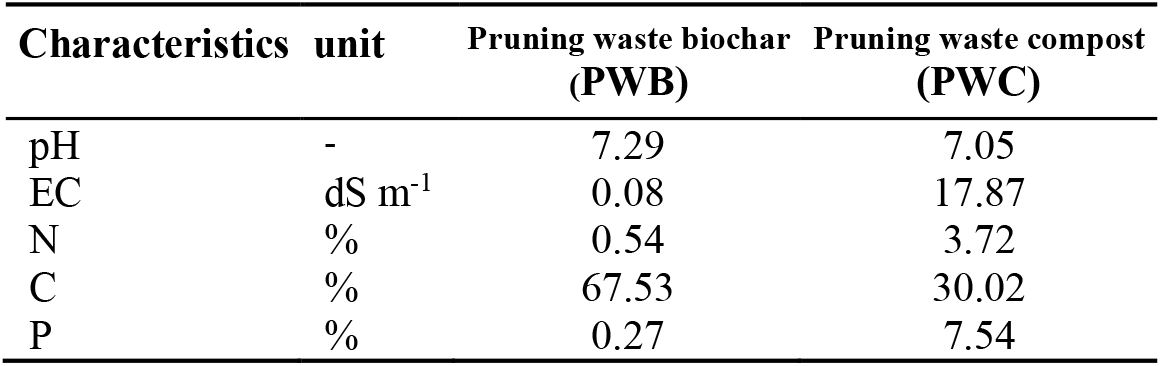
Characteristics of the tested PWC and PWB

| Characteristics | unit | Pruning waste biochar<br>(PWB) | Pruning waste compost<br>(PWC) |
| --- | --- | --- | --- |
| pH | - | 7.29 | 7.05 |
| EC | dS m<sup>-1</sup> | 0.08 | 17.87 |
| N | % | 0.54 | 3.72 |
| C | % | 67.53 | 30.02 |
| P | % | 0.27 | 7.54 |

### 2.3. Greenhouse Experiment

#### 2.3.1. Rhizobox experiment

Lab-built rhizoboxes with dimensions of 20 × 15 × 20 cm (length × width × height) were used in the greenhouse experiments. The inner space of the rhizoboxes was divided into two zones using 325 mesh nylon cloth: (i) a 2 cm thick rhizosphere zone, and (ii) a 5.8 cm thick non-rhizosphere zone on either side of the rhizosphere zone (Fig. 1). PWB and PWC derived from apple trees and grape vines was incorporated into the soil so as to provide 1.5% net organic C; each box thus contained 5.80 kg soil complemented with 41.19 g compost kg^−1^ soil (PWC), 22.21 g biochar kg^−1^ soil (PWB), or sterilized inoculated soil (control). In addition, a rock phosphate source providing 80 mg P kg^−1^ soil was placed 5 cm below the seeds to act as an insoluble P source.

**Figure 1.**
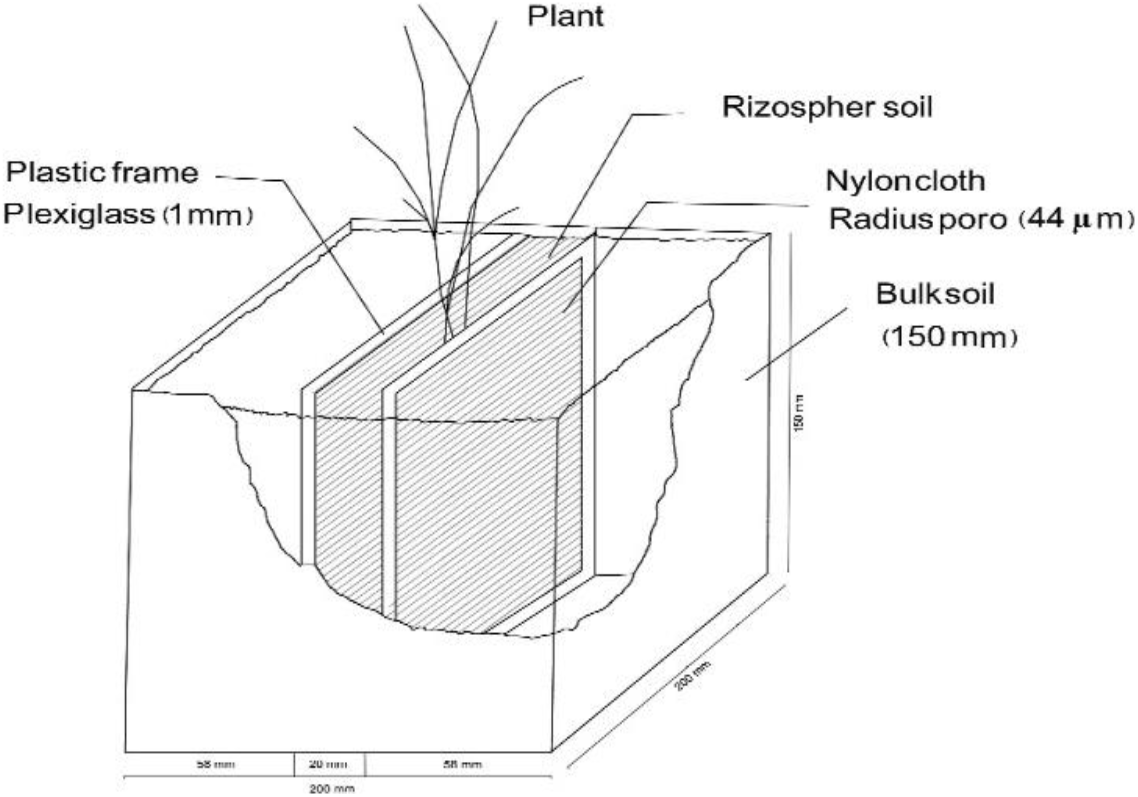
A schematic of rhizobox

Microbial inoculation was performed using microbial strains from the microbial bank of the Soil Science Department of Urmia University and included the mycorrhizal fungus *Glomus fasciulatum*. The amount of mycorrhiza inoculum was 70 g per box and it was uniformly dispersed under the seeds with a 0.5-cm spacing [27]. After adding the inoculum, seeds of the wheat cultivar ‘Pishtaz’ (*Triticum aestivum* L.) were disinfected with 0.5% sodium hypochlorite and sown in the rhizosphere zones of the rhizoboxes. After the seeds germinated, the seedlings were thinned to provide four plants per rhizobox. During their growth periods, the plants were irrigated with distilled water and fed with P-free Rorison nutrient solution to satisfy their nutrient requirements [28].

#### 2.3.2. Soil analysis

At the end of the 65-day wheat growth period, two soil samples were collected from the non-rhizosphere zone (specifically, from the 2 cm furthest from the rhizosphere zone) and combined. Rhizosphere soil was then sampled by removing the plexiglass framework of the box and then removing the soil from the box. Main and secondary roots were then carefully removed from the rhizosphere soil and two samples were collected and combined. Two final samples were thus obtained from each rhizobox, one representing the rhizosphere zone and the other representing the non-rhizosphere zone. The OC [24], MBC [29], MBP [30], BR [31], SIR [32], ACP and ALP [33] and arbuscular mycorrhizal fungi colonization percentage [34] were then determined for each sample.

#### 2.3.3. Statistical Analysis

Statistical analyses including analysis of variance and comparison of data means was performed using Duncan’s Multiple Range Test with a significance threshold of *p* < 0.05 using the SAS software package (Version 9.2).

## 3. Results

Comparisons of means for the interaction between organic sources and AMF inoculation (*p* < 0.001) revealed that treatments involving AMF inoculation increased organic C and soil biological parameters to a much greater degree than treatments without inoculation. Even without the addition of exogenous organic material, the soil biological parameters after inoculation were substantially higher than under non-inoculated control conditions. However, in the absence of added organic matter, AMF inoculation did not significantly increase the soil’s organic C content (Table 3). The highest soil organic C content and biological parameter values were observed for the treatment combining AMF inoculation with compost addition. However, the organic C content did not differ significantly between the compost and biochar treatments under non-inoculation conditions. Biochar treatments yielded higher organic C levels than control treatments under both inoculation and non-inoculation conditions, but the increase in organic C was lower than that caused by adding compost. A comparison of means for the interactive effect of organic source and microbial inoculation on MBC and MBP showed that the addition of organic matter and AMF inoculation had a significant effect (*p* < 0.001) when compared to the control treatment without AMF inoculation. In the compost treatments, the presence of AMF increased MBC and MBP levels 1.66- and 5.24-fold, respectively. Similarly, in the biochar treatments, AMF inoculation increased MBC and MBP levels 1.37- and 1.34-fold, respectively. Under non-inoculation conditions, adding compost increased MBC and MBP to a greater degree than adding biochar (Table 3). Compost treatment combined with AMF inoculation also increased BR and SIR, and this increase was 1.28 times greater in the presence of AMF than without inoculation. Similar trends were observed for biochar treatment.

**Table 3.** Effects of the organic amendment source and microbial inoculation on OC, MBC, MBP, BR and SIR

| Microbial inoculation | Organic sources | OC (%) | MBC (mg kg<sup>-1</sup>) | MBP (mg kg<sup>-1</sup>) | BR (mg CO<sub>2</sub> d<sup>-1</sup> kg<sup>-1</sup>) | SIR (mg CO<sub>2</sub> d<sup>-1</sup> kg<sup>-1</sup>) |
| --- | --- | --- | --- | --- | --- | --- |
| +AMF | PWB | 1.64<sup>b</sup> | 789.3<sup>b</sup> | 20.20<sup>b</sup> | 53.15<sup>b</sup> | 100.5<sup>b</sup> |
|  | PWC | 2.0<sup>a</sup> | 957.1<sup>a</sup> | 85.72<sup>a</sup> | 56.17<sup>a</sup> | 112.3<sup>a</sup> |
|  | Cont. | 0.44<sup>d</sup> | 429.0<sup>d</sup> | 7.43<sup>d</sup> | 25.97<sup>e</sup> | 51.99<sup>e</sup> |
| -AMF | PWB | 0.87<sup>c</sup> | 574.0<sup>c</sup> | 15.01<sup>c</sup> | 34.86<sup>d</sup> | 78.21<sup>d</sup> |
|  | PWC | 0.97<sup>c</sup> | 573.7<sup>c</sup> | 16.33<sup>c</sup> | 43.63<sup>c</sup> | 87.48<sup>c</sup> |
|  | Cont. | 0.32<sup>d</sup> | 264.5<sup>e</sup> | 3.52<sup>e</sup> | 8.61<sup>f</sup> | 15.46<sup>f</sup> |
| LSD<sub>0.05</sub> |  | 0.18 | 7.31 | 2.41 | 2.88 | 4.34 |

Means followed by the same letters are not significantly different according to Duncan's multiple range test at the $p < 0.05$ level. +AMF, -AMF, PWB, PWC, and Cont denote inoculation with arbuscular mycorrhizal fungi, without mycorrhizal inoculation, pruning waste biochar, pruning waste compost, and control (no added organic matter), respectively.

Adding organic matter (especially compost) to the soil caused the organic matter content of both the rhizosphere zone and the non-rhizosphere zone to differ significantly from that in control treatments (*p* < 0.05; Table 4). The soil organic carbon content was higher in the rhizosphere than in non-rhizosphere in all treatments, but the difference was not significant in all cases. The addition of organic matter to the rhizosphere and non-rhizosphere zones significantly increased their MBC and MBP contents (p <0.01) when compared to controls (Table 4). The highest MBC content was observed when rhizosphere soil was amended with compost; under these conditions, the MBC content of rhizosphere soil was 1.30% higher than that of non-rhizosphere soil. The MBC of the rhizosphere soil was lowest under organic matter-free control conditions; even under these conditions, the MBC of the rhizosphere soil was 3.49% higher than that of the non-rhizosphere soil. The highest MBP was observed in the rhizosphere zone after treatment with compost, which was 5.55% higher than the MBP in the non-rhizosphere. Biochar also improved these biological parameters when compared to the control, albeit to a lesser degree than compost; it increased both BR and SIR in the rhizosphere significantly more (*p* < 0.001) than in the non-rhizosphere zone (Table 4). The BR and SIR in both the rhizosphere and the non-rhizosphere were higher under the organic amendment treatments than under control conditions; the highest BR (5263 mg CO_2_ kg^−1^ soil) and SIR (103.3 mg CO_2_ kg^−1^ soil) were observed in rhizosphere zone after compost treatment.

**Table 4.** OC, MBC, MBP, BR and SIR in different soil zones with different organic amendments

| Soil | Organic sources | OC (%) | MBC (mg kg<sup>-1</sup>) | MBP (mg kg<sup>-1</sup>) | BR (mg CO<sub>2</sub> d<sup>-1</sup> kg<sup>-1</sup>) | SIR (mg CO<sub>2</sub> d<sup>-1</sup> kg<sup>-1</sup>) |
| --- | --- | --- | --- | --- | --- | --- |
| Rhizosphere | PWB | 1.29<sup>bc</sup> | 687<sup>c</sup> | 18.98<sup>c</sup> | 46.99<sup>b</sup> | 92.61<sup>b</sup> |
|  | PWC | 1.52<sup>a</sup> | 770.3<sup>a</sup> | 52.41<sup>a</sup> | 52.63<sup>a</sup> | 103.3<sup>a</sup> |
|  | Cont. | 0.42<sup>d</sup> | 352.7<sup>e</sup> | 6.71<sup>e</sup> | 20.27<sup>d</sup> | 37.13<sup>d</sup> |
| Non-rhizosphere | PWB | 1.22<sup>c</sup> | 676.3<sup>d</sup> | 16.23<sup>d</sup> | 41.03<sup>c</sup> | 86.06<sup>c</sup> |
|  | PWC | 1.45<sup>ab</sup> | 760.4<sup>b</sup> | 49.65<sup>b</sup> | 47.16<sup>b</sup> | 96.51<sup>b</sup> |
|  | Cont. | 0.35<sup>d</sup> | 340.8<sup>f</sup> | 4.24<sup>f</sup> | 14.31<sup>e</sup> | 30.33<sup>e</sup> |
| LSD<sub>0.05</sub> |  | 0.18 | 7.31 | 2.41 | 2.88 | 4.34 |

Means followed by the same letters are not significantly different according to Duncan's multiple range test at the p<0.05 level. PWB, PWC, and Cont. denote pruning waste biochar, pruning waste compost and Control (without organic matter), respectively.

AMF inoculation increased organic C in both the rhizosphere and non-rhizosphere zones when compared to the non-inoculated control treatment (Table 5). However, the organic C contents of the rhizosphere and non-rhizosphere did not differ significantly (*p* < 0.05) under either inoculation or non-inoculation conditions. Additionally, the microbial biomass C and P levels in the +AMF treatments were 1.22 and 3.25 times higher, respectively, than in the −AMF treatments (Table 5). On the other hand, MBC and MBP were much higher in the rhizosphere than in the non-rhizosphere (*p* < 0.001). AMF inoculation significantly (*p* < 0.001) increased BR and SIR in both the rhizosphere and non-rhizosphere zones when compared to controls (Table 5). Additionally, AMF inoculation increased BR and SIR in the rhizosphere by 13.06 and 7.95%, respectively, relative to the non-rhizosphere.

**Table 5.** OC, MBC, MBP, BR and SIR in rhizosphere and non-rhizosphere soil with and without microbial inoculation

| Microbial inoculation | Soil | OC (%) | MBC (mg kg<sup>-1</sup>) | MBP (mg kg<sup>-1</sup>) | BR (mg CO<sub>2</sub> d<sup>-1</sup> kg<sup>-1</sup>) | SIR (mg CO<sub>2</sub> d<sup>-1</sup> kg<sup>-1</sup>) |
| --- | --- | --- | --- | --- | --- | --- |
| +AMF | Rhizosphere | 1.39<sup>a</sup> | 730.9<sup>a</sup> | 39.08<sup>a</sup> | 47.86<sup>a</sup> | 91.64<sup>a</sup> |
|  | Non-rhizosphere | 1.33<sup>a</sup> | 719.4<sup>b</sup> | 36.49<sup>b</sup> | 42.33<sup>b</sup> | 84.89<sup>b</sup> |
| -AMF | Rhizosphere | 0.75<sup>b</sup> | 475.7<sup>c</sup> | 12.99<sup>c</sup> | 32.06<sup>c</sup> | 63.73<sup>c</sup> |
|  | Non-rhizosphere | 0.69<sup>b</sup> | 465.6<sup>d</sup> | 10.26<sup>d</sup> | 26.01<sup>d</sup> | 57.04<sup>d</sup> |
| LSD<sub>0.05</sub> |  | 0.18 | 7.31 | 2.41 | 2.88 | 4.34 |

Means followed by the same letters are not significantly different at the $p < 0.05$ level according to Duncan's test. +AMF and -AMF denote inoculation with arbuscular mycorrhizal fungi and without mycorrhiza, respectively.

The highest activity of phosphatase enzymes was measured in compost applied inoculated with AMF. AMF inoculation increased ACP and ALP enzymes by 5.92 and 5.01 times compared to non-inoculated control, respectively. The lowest activity of phosphatase enzymes was observed in non-inoculated control that was not amended with organic matter. Among non-inoculated organic treatments, compost showed higher activity of these enzymes than biochar, although biochar enhanced these enzymes’ activity compared to control. Also, its effect under AMF inoculation on the increased activity of the enzymes was in the second rank of magnitude after compost (Figure 2a). Organic treatments increased the activity of phosphatase enzymes in both rhizosphere and non-rhizosphere when compared to control (Figure 2b). According to the results, when PWC was applied, the activity of ACP and ALP enzyme was 1.06 and 1.04 times as great in the rhizosphere as in the non-rhizosphere region, and both regions showed significant differences when they were treated with compost than treated with biochar and control (Figure 2b). According to Figure 2b, it is observed that in all treatments, the activity of the ALP enzyme is greater than that of ACP. In control treatments, too, compost was more effective than biochar in prospering enzymatic activity. AMF inoculation increased the activity of phosphatase enzymes so that the activity of ACP and ALP enzymes under inoculation conditions was 1.30 and 2.07 times as great in the rhizosphere compared to non-rhizosphere region (Figure 2c).

**Figure 2.**
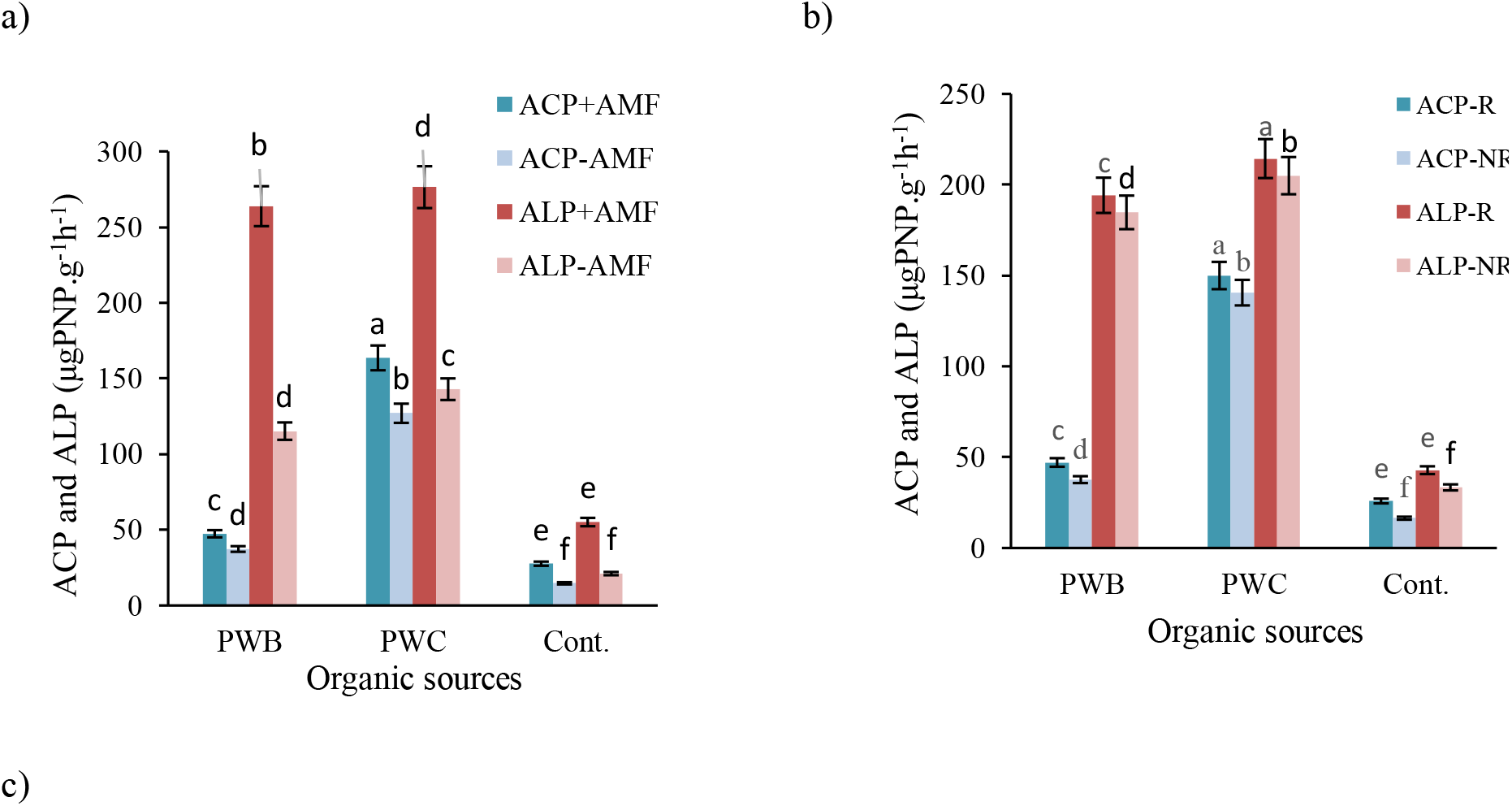
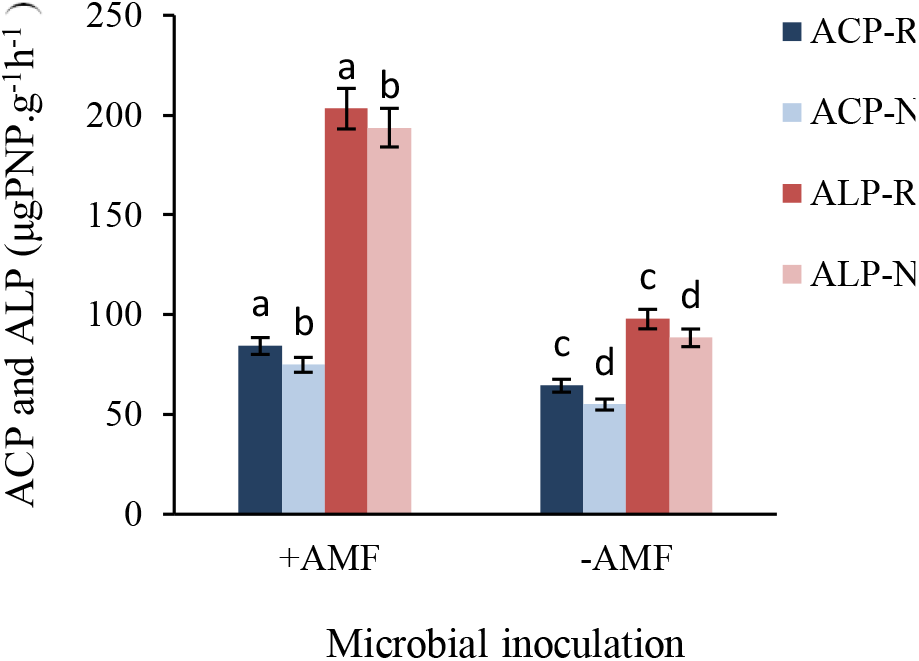
Means comparison of a) organic sources and microbial inoculation (+AMF, −AMF) on the soil ACP and ALP b) organic sources on the soil ACP and ALP, and c) microbial inoculation on the soil ACP and ALP. PWB, PWC, Cont., ACP, ALP, +AMF, −AMF, R and NR are pruning waste biochare, pruning waste compost, control (no organic mater), acid and alkaline phosphatase enzymes, inoculation with arbuscular mycorrhizal fungi, without mycorrhiza, rhizosphere and non-rhizosphere regions, respectively.

Finally, analysis of variance revealed that organic amendment and AMF significantly increased root mycorrhizal colonization (*p* < 0.001) when compared to controls (Fig. 3). Colonization was highest under the PWC and PWB treatments (51.45 and 41.24% higher than controls, respectively).

**Figure 3.**
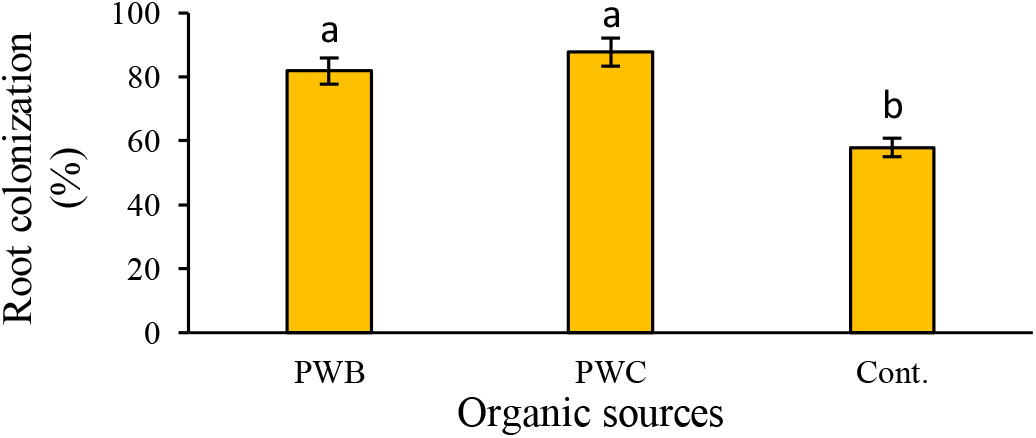
The effect of the organic source and AMF on root colonization (%) PWB, PWC and Cont. denote pruning waste biochare, pruning waste compost, and control (no organic mater), respectively.

## 4. Discussion

Biochar and compost amendment both increased the soil’s organic C content, but when organic amendment was combined with AMF inoculation, amendment with compost increased organic C more than amendment with biochar. This may be due to the stable carbon skeleton of biochar, which is likely to resist microbial decomposition and may thus enable a slow gradual release of organic C into the soil. For example, there is evidence that the lifespan of biochar carbon in soil is between 100 and 1000 years, which is 10-1000 times longer than that of soil organic carbon, making it a valuable long-term carbon source [35]. The increase in MBC following organic amendment may be due to the introduction of beneficial substrates for microorganisms that stimulate their activity and increase soil biological activity [36]. Accordingly, Babalol et al. reported that applying plant residue compost to soil increased its population of microorganisms, leading to an increase in MBP [37]. Amendment with wood biochar can also increase soil microbial activity both by providing a favorable habitat for microorganisms and by serving as a source of moisture, carbon, labile resources, and nutrients. Fig. 4 presents scanning electron microscopy (SEM) images of the apple and grape biochars used in this work, revealing that they consist of irregularly shaped porous particles. The porosity of biochar gives it a high specific area, enabling the adsorption of large quantities of dissolved organic matter, gases, and minerals. Additionally, the diameter of the pores in wood-derived biochars ranges from 2 to 80 μm; this size range is adequate to support mycorrhizal fungi [38]. Consequently, biochar and provides a very favorable habitat for microorganisms, especially AMF; its pores provide shelter against predators and protection from drought while also hosting resources that help satisfy their carbon, energy, and nutrient requirements. Consequently, biochar amendment increases the biological activity of mycorrhizal fungi [13]. Jin reported that the interaction of biochar with AMF increased the population of mycorrhizal fungi and the MBC content of rhizospheric soil to a greater degree than in non-rhizosphere soil [39]. The increase in MBC was more pronounced among mycorrhizal fungi than bacteria, which was attributed to more efficient use of resources by the fungi and more effective carbon assimilation due to the extensive hyphal network of the fungi. Importantly, fungi can establish hyphal bridges between biochar and plant roots.

**Figure 4.**
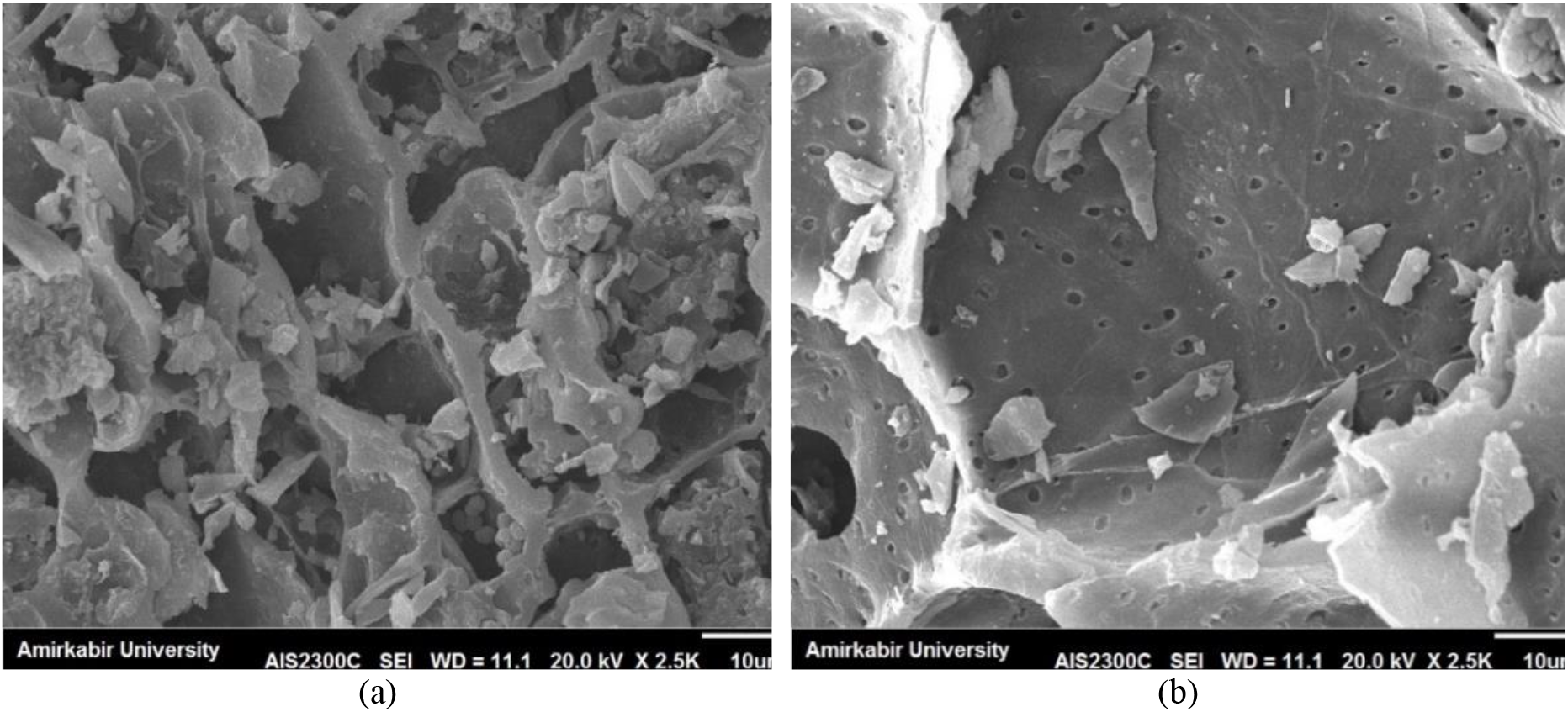
Scanning electron microscope (SEM) imagery of biochar of (a) apple pruning waste and (b) grape pruning waste at pyrolysis temperature of 350°C

Amendment with compost increases the organic C content of the soil as well as both plant growth and microbial activity. This normally leads to increased respiration (both BR and SIR). Compost amendment induced a greater increase in both rates of respiration than amendment with biochar, which may be related to its greater ease of decomposition. Obviously, the availability of adequate nutrients is a key determinant of the activity and growth of microorganisms. The effect of biochar on respiration differs from that of compost due to its structure and long half-life; Liu et al. found that biochars did not significantly influence soil respiration [11], whereas Steiner et al. observed increased microbial activity and growth following biochar amendment [40]. In this work, despite an increase in the microbial population following the addition of glucose to the biochar-treated soil, the rate of soil respiration (SIR) was unchanged. These results imply that the application of nutrient-rich but slowly decomposing biochars can support microbial population growth. Biochars also had some positive impacts on soil biological characteristics; the formation of biochars from wood at low pyrolysis temperatures leads to the retention of diverse compounds that promote microbial growth (e.g., sugars and aldehydes) on the surface of the biochar [41]. The short-term increase in microbial activity and population as well as soil respiration following biochar amendment is probably related to these labile constituents of freshly prepared biochars [42].

In general, the rhizosphere is the main zone in which organic compounds accumulate and decompose. It is thus unsurprising that it contains more organic matter than the non-rhizosphere. It seems that the active carbon of compost is partially decomposed after the compost is applied and that some of this carbon is added to the soil’s carbon reserve, increasing the organic carbon content of the soil. Biochar amendment also increased the soil’s organic carbon content, but to a lesser degree than treatment with compost. Zaman et al. reported that the application of urban waste compost increased soil MBC by enhancing the soil’s content of N, C, and soluble organic C [43]. This is consistent with our finding that the soil’s organic carbon content (Table 4) and total absorbed nitrogen (Table 2) were higher after compost amendment than after treatment with other organic materials. Additionally, De Neergaard and Magid found that the application of organic matter improved the MBC content in the rhizosphere of rye to a greater degree than that of the non-rhizosphere zone [44], and a study on the response of the soil microbial community to biochar derived from wheat and willow trees, Watzinger et al. indicated that biochar application increased microbial biomass and changed the microbial community structure [45]. MBP can be either a vital sink for soluble P when microbes compete with plants for P or a crucial source of P that helps partially meet plants’ P requirements. Soil microbes can thus help maintain stable levels of labile P in soils with low P availability. Redel et al. incorporated compost into soil and observed a positive association between MBP content and Olsen P (determined by extraction with sodium bicarbonate) [46]. Similarly, in this work, compost amendment increased available P in the soil more than any other treatment (Table 2). Biochar amendment has also been reported to increase MBP, possibly due to its high C content and positive effect on nutrient availability. Compost amendment strongly increased the microbial respiration rate because compost is rich in oxygenated organic compounds such as carbohydrates that are more readily degraded by microbes than aromatic carbon and alkyl carbon chains [47]. However, an earlier study on the effect of biochar on BR under rhizobox conditions [48] found that biochar increased BR, leading to a significantly higher rate of microbial respiration in the rhizosphere than in the non-rhizosphere zone.

Microorganisms can be regarded as active organic particles in the soil with charged and catalytically active surfaces and the ability to produce and exude a wide range of organic compounds including carbohydrates, and enzymes [49]. A soil’s MBC content is widely interpreted as a measure of its content of soil microbes and their activity. However, it does not distinguish between living and dead microbes; it is likely that AMF cells are included in the soil microbial biomass when their life cycle ends. The increase in MBC following AMF inoculation may thus be partly due to the addition of dead cells to the rhizosphere soil, as suggested in earlier studies on the maize rhizosphere [50]. The beneficial effects of the rhizosphere on the soil biological characteristics have also been highlighted by Zhao et al. [51]. In addition to the increases in rhizospheric MBC due to organic matter amendment and microbial inoculation, plants release organic compounds including sugars, amino acids, and vitamins into the rhizosphere, further reinforcing its differences from the non-rhizosphere zone. These root exudations also increase microbial biomass by making the rhizosphere more hospitable to microbes, which in turn increases plant growth and the production of microbial metabolites [52]. Consequently, the rhizosphere is characterized by elevated microbial activity and has a higher content of organic matter and nutrients than the rest of the soil, allowing it to play a key role in meeting plants’ nutrient requirements to support photosynthesis while also supporting a diverse microbial community leading to high levels of soil biological activity (and thus high levels of BR, SIR, MBC, and MBP) [53]. Organic matter application can boost microbial activity to improve enzymatic activities via three ways: (i) the application of organic matter to soil enhances soil microbial activity by microorganism-based energy pathway, resulting in enzyme synthesis; (ii) the enzymes accompanying organic matter reduce energy for the activity of microorganisms of soil for the synthesis of the enzymes, thereby increasing their activity and the rate of organic matter decomposition in soil; (iii) organic matter is partially decomposed in soil and then, it protects the enzymes against enzymatic hydrolysis by adsorbing and physically grasping the enzymes [54]. The enhancement of the activity of soil phosphatase enzymes under the treatments with PWC and PWB may be associated with the increased level of microbial biomass in response to the applied organic matter, soil nutrients, and the improvement of soil (physical, chemical and biological) characteristics, like nutrient and water holding capacity. Also, the higher performance of PWC in enhancing the activity of phosphatase enzymes as compared to PWB is likely to reflect its higher rate of decomposition and the availability of its N content, which stimulates microbial activity. The decline of ACP enzyme activity versus ALP under the application of organic matter, especially biochar, was not surprising because the studied soil, which had alkaline pH and calcareous conditions (Table 1), and the applied biochar, which had alkaline pH (Table 2), were not suitable for the activity of this enzyme. Jin et al. reported that the activity of ACP enzyme was decreased after the application of manure biochar, whereas the activity of ALP enzyme was increased [55]. The loss of ACP enzyme activity was attributed to the increase in soil pH by biochar application. Fungi need extra-cellular enzymes, such as phosphatase, to decompose the substrate of their surrounding medium into smaller molecules and to mobilize these molecules into the cells and also to use them in their metabolic activities. Thus, the medium around a fungus is studied with the exudation of extra-cellular enzymes. Since extra-cellular enzymes decompose promptly, therefore they should be adsorbed to soil particles to maintain their activities. In this respect, specific areas are of particular importance. These areas can be the surface of soil particulates, plant root, or organic matter like compost or biochar. The activity of extra-cellular enzymes depends on the site of the enzyme that is interacting with the surface of biochar particles [56]. If the enzyme’s active site is not covered and is operational and free to react with environment, its activity will be augmented. Still, if it is blocked, its activity will decline [57]. Therefore, some groups of enzymes become more active by the application of biochar and the other groups lose their activity depending on molecular composition, folding properties, and their adsorption potential [58]. The augmented activity of phosphatase in the rhizosphere versus non-rhizosphere can be associated with the microbial and root activity of the plant. Similarly, in a study on the effect of organic fertilizer on the activities of phosphatase enzymes in the rhizosphere of plants using rhizobox, Balik et al. reported that the action of ACP and ALP enzymes were significantly increased in bulk soil [59].

The rate by which microorganisms can synthesize and release phosphatase varies with soil pH. As pH is increased, ALP becomes more stable and active so that its activity is maximized at pH = 11. In addition, ACP is mostly exuded by fungi and the optimal pH for it is acidic to neutral conditions. Still, bacteria predominantly exude ALP and are more active at pH > [60]. Furthermore, ACP is produced mainly by plants and microorganisms, but microorganisms predominantly exude ALP.

Organic matter amendment also improves the physical properties of the soil, making it better able to support the development and activity of microorganisms and various bioprocesses [61]. The improvements in soil biochemical properties and nutrient availability induced by PWC or PWB amendment were reinforced by inoculation with AMF, and also enhanced root mycorrhizal colonization. Similarly, Mäder et al. reported that the application of compost to soil increased mycorrhizal colonization by 30-60% in the wheat rhizosphere [62]. Amendment with hardwood biochar (2% w/w) similarly increased root colonization, which was attributed to increased growth of the external hyphae of arbuscular mycorrhiza [17]. Biochar can influence the symbiosis of arbuscular mycorrhiza and plant roots by (i) changing the availability of nutrients affecting the relationships between plants and AMF and/or their physicochemical properties, (ii) stimulating microbial activity and population growth by strengthening mycorrhizal symbiosis, (iii) disturbing chemical signals or detoxifying chemicals that inhibit the activity of mycorrhiza fungi, and (iv) providing physical shelter for mycorrhiza fungi against adverse conditions.

## 5. Conclusion

The results presented herein show that combining organic matter amendment with microbial inoculation can have significant positive effects on soil quality parameters. Adding organic matter such as compost or biochar to the soil and inoculation with AMF in the root zone strongly increased soil biological activity, with particularly strong increases in soil quality parameters (BR, SIR, MBC, and MBP) occurring in the rhizosphere. Combined compost amendment and AMF inoculation led to the highest OC, MBC, and MBP values, but the effect of biochar amendment on root mycorrhizal colonization was similar to that of adding compost. In general, amendment with compost had a stronger positive effect on the studied soil biological parameters than treatment with biochar, which may be due to the structure and greater stability of biochar. It should be noted that not all soil quality parameters respond in the same way to microbial inoculation and organic matter amendment, and that the efficiency of plant responses to these treatments will depend on the experimental conditions including the plant type and species, the experiment type (pot or field study), the choice of organic matter, and the nutrient ratios. Consequently, it will be necessary to confirm the results presented herein by studying the combined effects of organic amendment and microbial inoculation in greenhouse and field studies on various plant species and to assess the economic viability of organic amendment, especially with biochar. Nevertheless, the results presented herein show that combining organic matter amendment with AMF inoculation can strongly increase the quality, biological activity, and biodiversity of calcareous alkaline and may represent a very promising alternative to the costly application of chemical fertilizers.

## Author Contributions

R.V. guided *g*reenhouse experiments and took part in drafting the manuscript. M.H.R.S. conceived and supervised the work. M.B. provided guidance and manuscript reviews. R.R.V. reviewed and edited the manuscript. All authors have read and agreed to the published version of the manuscript.

## Institutional Review Board Statement

Not applicable.

## Informed Consent Statement

Not applicable.

## Data Availability Statement

The data presented in this study are available on request from the corresponding author.

